# Why cooperation is not running away

**DOI:** 10.1101/316117

**Authors:** Félix Geoffroy, Nicolas Baumard, Jean-Baptiste André

**Affiliations:** ISEM, Univ. Montpellier, CNRS, IRD, EPHE, Montpellier, France; Institut Jean Nicod, Département d’études cognitives, ENS, EHESS, CNRS, PSL Research University, Paris, France

**Keywords:** partner choice, biological markets, matching models, competitive altruism, human cooperation

## Abstract

A growing number of experimental and theoretical studies show the importance of partner choice as a mechanism to promote the evolution of cooperation, especially in humans. In this paper, we focus on the question of the precise quantitative level of cooperation that should evolve under this mechanism. When individuals compete to be chosen by others, their level of investment in cooperation evolves towards higher values, a process called competitive altruism, or runaway cooperation. Using a classic adaptive dynamics model, we first show that, when the cost of changing partner is low, this runaway process can lead to a profitless escalation of cooperation. In the extreme, when partner choice is entirely frictionless, cooperation even increases up to a level where its cost entirely cancels out its benefit. That is, at evolutionary equilibrium, individuals gain the same payoff than if they had not cooperated at all. Second, importing models from matching theory in economics we, however, show that, when individuals can plastically modulate their choosiness in function of their own cooperation level, partner choice stops being a runaway competition to outbid others, and becomes a competition to form the most optimal partnerships. In this case, when the cost of changing partner tends toward zero partner choice leads to the evolution of the socially optimum level of cooperation. This last result could explain the observation that human cooperation seems to be often constrained by considerations of social efficiency.

## 1 Introduction

Cooperation among non-kin constitutes a puzzle for evolutionary biologists, and a large body of theoretical models, inspired by game theory, have been developed to solve it. The most commonly accepted explanation is that cooperation can be enforced if it triggers a conditional response on the part of others [102]. Several enforcement mechanisms have been proposed: direct reciprocity [15, 68, 98], indirect reciprocity [70, 77, 78], punishment [30, 32, 33] and partner choice [35, 75, 76, 84]. A growing number of experimental studies support the idea that, among this set of mechanisms, partner choice is likely to be particularly influential in nature, both in inter-specific and in intra-specific interactions [34, 52, 59, 64, 65, 86, 90]. Besides, partner choice is also believed to play a major role in human cooperation, where friendships and coalitions are common [16, 19, 22] (see also Discussion).

The key idea of partner choice models is that, when one happens to be paired with a defecting partner, one has the option to seek for another, more cooperative, partner present in the “biological market” and interact with her instead of the defector. This possibility allows cooperators to preferentially interact with each other, and, consequently, prevents any invasion by free-riders [3, 4, 17, 25, 35, 48, 49, 53, 75, 76, 84].

So far, the primary objective of most partner choice models has been to explain how *some* cooperation can exist at all in an evolutionary equilibrium. On this ground, models have reached a clear answer: partner choice can trigger the evolution of cooperation. In this paper, however, we are interested in another issue that models generally consider with less scrutiny: that of understanding the quantitative *level* of cooperation that should evolve under partner choice.

This analysis is crucial because the quantitative level of cooperation determines the “social efficiency”, also called the Pareto efficiency, of interactions. Cooperating too little is inefficient because individuals miss some opportunities to generate social benefits. But cooperation, as any investment, is likely to have diminishing returns [8, 66, 101]. As a result, there is a “socially optimal” amount of cooperation, an intermediate level where the sum of the helper and helpee’s payoff is maximized. Cooperating more than this amount is hence also inefficient, because it increases more the cost of cooperation than it raises its benefit. In the extreme, there is even a “wasteful” “wasteful” threshold beyond which the overall cost of cooperation becomes larger than its benefit. If two partners cooperate more than this threshold, the net benefit of their interaction is negative, that is they are both worst off than if they had not cooperated at all.

Prima facie, partner choice appears to be a unidirectional pressure acting on the evolution of cooperation, unlikely to generate an intermediate equilibrium. Competition to be chosen by others, called “competitive altruism” [61, 74, 80], should lead to a runaway of cooperation, as it does in sexual selection [104]. In principle, this runaway should proceed up to the point where the cost of investing into cooperation cancels out the benefit of finding a partner [50, p. 152, 103] that is up to the “wasteful” threshold where cooperation becomes fruitless. Is competitive altruism, however, balanced by opposite forces, leading to an evolutionary stabilization of cooperation below this threshold? Is this level socially optimal, or does partner choice lead to the investment into counterproductive forms of cooperation to out-compete others as it does in sexual selection?

In the theoretical literature on partner choice, relatively little attention has been given to these questions. First of all, a large proportion of models consider cooperation as an all-or-nothing decision and thus cannot study its quantitative level [4, 5, 25, 37, 39, 40, 48, 53, 62, 89, 94, 108]. Second, some models consider cooperation as a quantitative trait but do not entail diminishing returns, and are thus ill-suited to study the social efficiency of cooperative interactions [51, 74, 88, 92]. Third, still other models consider cooperation as a quantitative trait with diminishing returns, but they only focus on one side of the problem -the evolution of cooperation- considering the other side -the strategy employed by individuals to choose their partner– as an exogenous parameter [17, 49, 105, 106].

To our knowledge, only one existing model studies the joint evolution of cooperation and partner choice in a quantitative setting with diminishing returns (McNamara *et al.* 2008) [72]. However, McNamara *et al.* [72] make two key assumptions that turn out to have important consequences: (i) they assume that variability in the amount of cooperation is maintained owing to a very large genetic mutation rate on this trait, which prevents natural selection to act efficiently, and (ii) they restrict the set of possible strategies to choose one’s partner in such a way that individuals can never do so in an optimal manner.

In this paper, we build a model inspired by McNamara *et al.* [72], in which a quantitative level of cooperation expressed by individuals jointly evolves with a quantitative level of choosiness regarding others’ cooperation, while relaxing these two assumptions. First, we observe that competition to be chosen as a partner leads to a joint rise of both cooperation and choosiness up to a level that depends on the efficiency of partner choice that is, in particular, on the cost of changing partner. The more efficient is partner choice, the higher cooperation is at evolutionary stability. Moreover, when the cost of changing partner is low, cooperation can rise beyond its socially optimal level. In fact, in the limit where partner choice is entirely frictionless (i.e. the cost of changing partner is zero), cooperation and choosiness rise up to the “wasteful threshold” where the cost of cooperation entirely cancels out its benefit. Individuals gain the same payoff than if they had not cooperated at all. Hence, at first sight, our analyses show that partner choice generates no systematic trend toward the socially optimal level of cooperation.

However, we then import tools from the economics literature and assume that individuals can plastically modulate their choosiness in function of their own cooperation level. This plasticity allows every individual to behave optimally on the biological market, which did not occur in the first model. In this second approach, we show that assortative matching emerges. That is, more cooperative individuals are also choosier and thus interact with more cooperative partners. As a consequence of this assortment, and provided that partner choice is efficient enough, cooperation evolves to the socially optimal level, where the mutual efficiency of cooperation is maximised.

## 2 Methods

### 2.1 Partner choice framework

We model partner choice in an infinite size population using Debove *et al.*’s framework [43]. Solitary individuals randomly encounter each other in pairs at a fixed rate *β*. In each encounter, the two players decide whether they accept one another as a partner (see below how this decision is made). If one of the two individuals (or both) refuses the interaction, the two individuals immediately split and move back to the solitary pool. If both individuals accept each other, on the other hand, the interaction takes place and lasts for an exponentially distributed duration with stopping rate *τ*, after which the two individuals move back to the solitary pool again. The ratio *β/τ* thus characterizes the “fluidity” of the biological market. If *β* is high and *τ* is low, individuals meet each other frequently and interact for a long time. In such an almost frictionless market, partner choice is almost cost-free so they should be choosy about their partner’s investment in cooperation. Conversely, if *β/τ* is low, individuals rarely meet potential partners and interact for a short time. In such a market, on the contrary, individuals should accept any partner.

Regarding the encounter rate, here we assume that *β* is a fixed constant independent of the density of available partners, an assumption called “linear search” that captures a situation in which already paired individuals do not hinder the encounters of solitary individuals [46]. In the Supplementary Information, however, using simulations we also analyse the model under the assumption that *β* increases linearly with the proportion of solitary individuals in the population, an assumption called “quadratic search” that corresponds to a situation in which already matched individuals interfere with the encounters of solitary individuals (and that is also equivalent to the classic mass-action kinetics used in mathematical epidemiology). In the paper, we only describe the results obtained under linear search. The results obtained under quadratic search are qualitatively similar (see the Supplementary Information).

Regarding the nature of the social interaction, we consider a quantitative version of the prisoner’s dilemma in continuous time. Each individual *i* is genetically characterized by two traits: her cooperation level *x*_*i*_, and her choosiness *y*_*i*_. Cooperation level *x*_*i*_ represents the quantitative amount of effort that an individual *i* is willing to invest into cooperation. Choosiness *y*_*i*_ represents the minimal cooperation level that an individual *i* is willing to accept in a partner, i.e. every potential partner *j* with cooperation *x*_*j*_ ≥ *y*_*i*_ will be accepted, whereas every potential partner with *x*_*j*_ < *y*_*i*_ will be rejected. Once an interaction is accepted by both players, at every instant of the interaction, each player invests her effort *x*_*i*_ (see below for the payoff function), and the interaction lasts in expectation for 1/*τ* units of time, where *τ* is the stopping rate of the interaction.

When they are solitary, individuals gain a payoff normalized to zero per unit of time. When involved into an interaction, they gain a social payoff that depends on both partners’ cooperation level. The cooperative interaction is a continuous prisoner’s dilemma: making an investment brings benefits to the partner but comes at a cost to the provider. As stated in the introduction, we make the additional assumption that cooperation has diminishing returns [8, 66, 101]. This induces the existence of an intermediate level of cooperation at which the sum of the partners’ gains is maximized, the so-called “social optimum”. An individual *i* paired with *j* gains the following social payoff Π(*x*_*i*_, *x*_*j*_) per unit of time:

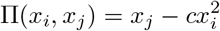

Hence, the expected payoff of an individual *i* paired with *j* is

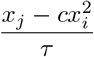

where *τ* is the stopping rate of the interaction. The socially optimal level of cooperation is 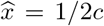. Beyond this level, the net benefit of cooperation decreases. Eventually, the interaction becomes entirely profitless, or even costly, if individuals invest more than the “wasteful threshold” *x* = 1/*c*. We allow both cooperation and choosiness to take any positive real value.

Previous studies demonstrated that the existence of some variability among individuals is necessary to stabilize conditional cooperation [49, 51, 72, 73, 92]. If every possible partner is equally cooperative, then there is no need to be choosy with regard to the quality of one’s partner, and choosiness cannot be evolutionarily stable. In order to capture the effect of variability in the simplest possible way, we assume that individuals do not perfectly control their investment into cooperation (as in Song & Feldman [92] and André [9] for instance). An individual’s actual cooperation level *x*_*i*_ is a random variable which follows a truncated to zero normal distribution around the individual’s gene value 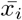, with standard deviation *σ*. In what follows, we call cooperation level the genetically encoded cooperation level that individuals aim for, and “phenotypic cooperation” the actual level of cooperation that they express after phenotypic noise. For the sake of simplicity, here, we assume that an individual’s cooperation level is randomized at every encounter. In the Supplementary Information, however, we also consider the alternative assumption where phenotypic noise occurs only once at birth (see also section 3.1).

We are interested in the joint evolution of cooperation, and choosiness by natural selection. We undertake and compare the consequences of two distinct assumptions. In a first approach, we assume that both cooperation and choosiness are hard-wired traits, that is each individual is characterized by a single level of cooperation 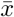 and a single choosiness *y*, both expressed unconditionally. In a second approach, we still assume that cooperation is a hard-wired trait, but we consider that choosiness is a reaction norm by which individuals respond to their own phenotypic cooperation.

### 2.2 Hard-wired choosiness

Here, we assume that each individual is genetically characterized by two traits: his level of cooperation 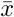 and his choosiness *y* and we are interested in the evolution of these two traits by natural selection. For this, we need to derive the fecundity of a rare mutant m playing strategy 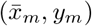 in a resident population *r* playing strategy 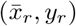. The mutant fecundity is proportional to her cumulative lifetime payoff *G*_*m*_, which can be written as (see SI for a detailed analysis of the model):

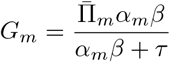

with *α*_*m*_ the mean probability for an encounter between the mutant and a resident to be mutually accepted, and 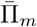 the mutant mean social payoff (see Table 1 for a list of the parameters of the model). This expression is similar to the classical sequential encounter model of optimal diet [87].

The evolutionary trajectory of the two traits (choosiness and cooperation) can be studied from the analysis of the selection gradient on each trait:

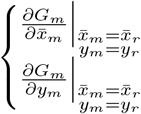

We could not derive an analytical expression of the evolutionarily stable strategy. However, we numerically computed the selection gradient on each trait, in order to study the evolutionary trajectories.

**Table 1.** Parameters of the model

| Parameter | Definition |
| --- | --- |
| $\bar{x}_i$ | Cooperation level of individual $i$ (mean value before applying noise) |
| $y_i$ | Choosiness of individual $i$ |
| $\sigma$ | Standard deviation of the phenotypic cooperation distribution |
| $\beta$ | Encounter rate |
| $\tau$ | Split rate |
| $\Pi(x_i, x_j)$ | Social payoff of an individual $i$ matched with a partner $j$ |
| $c$ | Cost of cooperation |
| $\alpha_i$ | Mean probability for an individual $i$ to interact when she encounters a resident |
| $\bar{\Pi}_i$ | Mean social payoff for an individual $i$ interacting with a resident |
| $G_i$ | Cumulative lifetime payoff of an individual $i$ |

### 2.3 Plastic choosiness

Because cooperation is subject to phenotypic noise (i.e. one does not perfectly control one’s own level of cooperation), it could make sense, at least in principle, for individuals to adapt plastically their degree of choosiness to the actual phenotypic cooperation that they happen to express. For instance, it could make sense for those individuals who happen to be phenotypically more generous to be also choosier, and vice versa. In our second model, we aim to explore the consequences of this possibility. To do so, we assume that choosiness is not a hard-wired trait, but a plastic decision that individuals take in function of their own phenotypic cooperation. An individual’s “choosiness strategy” is thus defined as a reaction norm rather than a single value.

Our aim in this second model is to study the joint evolution of cooperation 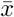 on one hand, and of the “choosiness strategy” *y*(*x*), defined as the shape of a reaction norm, on the other hand. One facet of this problem, therefore, consists in seeking for the equilibrium choosiness strategy in a situation where both one’s own quality (one’s phenotypic cooperation level) and the quality of one’s prospective partners vary. Matching theory, a branch of micro-economics, provides tools to resolve this problem. Here we briefly explain this approach, and show how it applies to our problem.

In a first category of approaches, called matching models, changing partner is assumed to be entirely cost-free [23, 55]. That is to say, agents have an infinite amount of time available to find each other. In this setting, theory shows that there is a unique equilibrium choosiness strategy: an individual with phenotypic cooperation *x* should only accept to interact with individuals with at least the same phenotypic cooperation level *x*, i.e. the equilibrium reaction norm is the identity function. This equilibrium strategy leads to a strictly positive assortative matching in which individuals are paired with likes.

The second category of approaches, called search and matching models, accounts for frictions in the matching process, i.e. incorporates an explicit cost for changing partner [38]. These models actually correspond exactly to our own partner choice framework. Individuals randomly encounter each other at a given rate and, when an individual refuses an interaction, she has to wait for some time before encountering a new partner. Unfortunately, the equilibrium choosiness reaction norm *y*\*(*x*) cannot be analytically derived in these models. However, Smith [91] has shown that a mathematical property of the social payoff function Π(*x*_*i*_, *x*_*j*_) allows predicting the shape of this reaction norm. If the social payoff function Π(*x*_*i*_, *x*_*j*_) is strictly log-supermodular, then *y*\*(*x*) is strictly increasing with *x*. If this is the case, the more an individual invests into cooperation, the choosier she should be. This equilibrium is called a weakly positive assortative matching. Log-supermodularity is defined as the following: Π(*x*_*i*_, *x*_*j*_) is strictly log-supermodular only if Π(*x*_*i*_, *x*_*j*_)Π(*x*_*k*_, *x*_*l*_) > Π(*x*_*i*_, *x*_*l*_)Π(*x*_*k*_, *x*_*j*_) for any investments *x*_*i*_ > *x*_*k*_ and *x*_*j*_ > *x*_*l*_.

Matching and search and matching models are, however, only interested in characterizing the equilibrium choosiness strategy of individuals, assuming a given, fixed, distribution of cooperation levels. As a result, matching models can offer an insight into the evolution of choosiness, but not into the joint evolution of choosiness and cooperation. To study this joint evolution in the case where choosiness is a reaction norm, and not a single value, we developed individual-based simulations.

### 2.4 Individual-based simulations

In addition to our analytical models, we run individual-based simulations coded into Python. We simulate the joint evolution of cooperation and choosiness in a Wright–Fisher population of *N* individuals, with the same lifespan *L* and non-overlapping generations. Mutations occur at rate *μ* and mutant genes are drawn from a normal distribution around the parent’s gene value, with standard deviation *σ*_*mut*_. Large effect mutations are implemented with probability *μ*_*l*_. They do not alter the equilibrium result and they allow to speed up the joint evolution process. We run long enough simulations for both choosiness and cooperation to stabilize. In contrast with previous papers [51, 73, 88], here we consider a continuous rather than discrete trait space, because Sherratt & Roberts [88] have shown than too much discretization can produce undesirable consequences when studying a joint evolution process. In the Supplementary Information, we also present additional simulations based on a Moran process with overlapping generations, where the lifespan of individuals is determined by a constant mortality rate (see also section 3.1 and [72]).

We run simulations both under the assumption that choosiness is hard-wired, and under the assumption that it is a reaction norm. In the second case, we test two types of reaction norms. First, we consider polynomial functions, the coefficients of which evolve by natural selection. Second, we consider step functions with evolving coefficients coding for the value of choosiness for each interval of cooperation. In the initial generation, all reaction norms are set to a constant zero function, so that individuals are never choosy at initiation.

## 3 Results

### 3.1 Hard-wired choosiness

Without variability in cooperation (*σ* = 0), there is no selective pressure to be choosier and, therefore, to be more cooperative. The only Nash equilibrium is 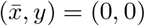, see SI for a demonstration.

When phenotypic cooperation is variable, however, the evolutionarily stable strategy cannot be formally derived. We therefore study the joint evolutionary dynamics of cooperation and choosiness by plotting numerically the selection gradients acting on both traits. In Figure 1, we show the evolutionary dynamics of cooperation, choosiness, and average payoff, in a case where partner choice is very effective. When starting from an initially selfish population, cooperation and choosiness jointly rise above zero (Fig. 1a). At first, this leads to an increase of the net social payoff (Fig. 1b) because cooperation is efficient (that is, the marginal benefit of increasing cooperation for the helpee is larger than its marginal cost for the helper). At some point, however, cooperation reaches the socially optimal level where the net payoff of individuals is maximized. Beyond this level, the marginal cost of increasing cooperation is larger than the marginal benefit, but the evolutionary runaway of cooperation and choosiness does not stop. Cooperation keeps on rising toward higher values, thereby decreasing the net payoff (Fig. 1b). Eventually, cooperation and choosiness stabilize when cooperation is so high, and therefore so inefficient, that its cost entirely cancels out its benefit (the so-called “wasteful threshold”). That is, at ESS, individuals gain the same payoff than if they had not cooperated at all.

**Figure 1.**
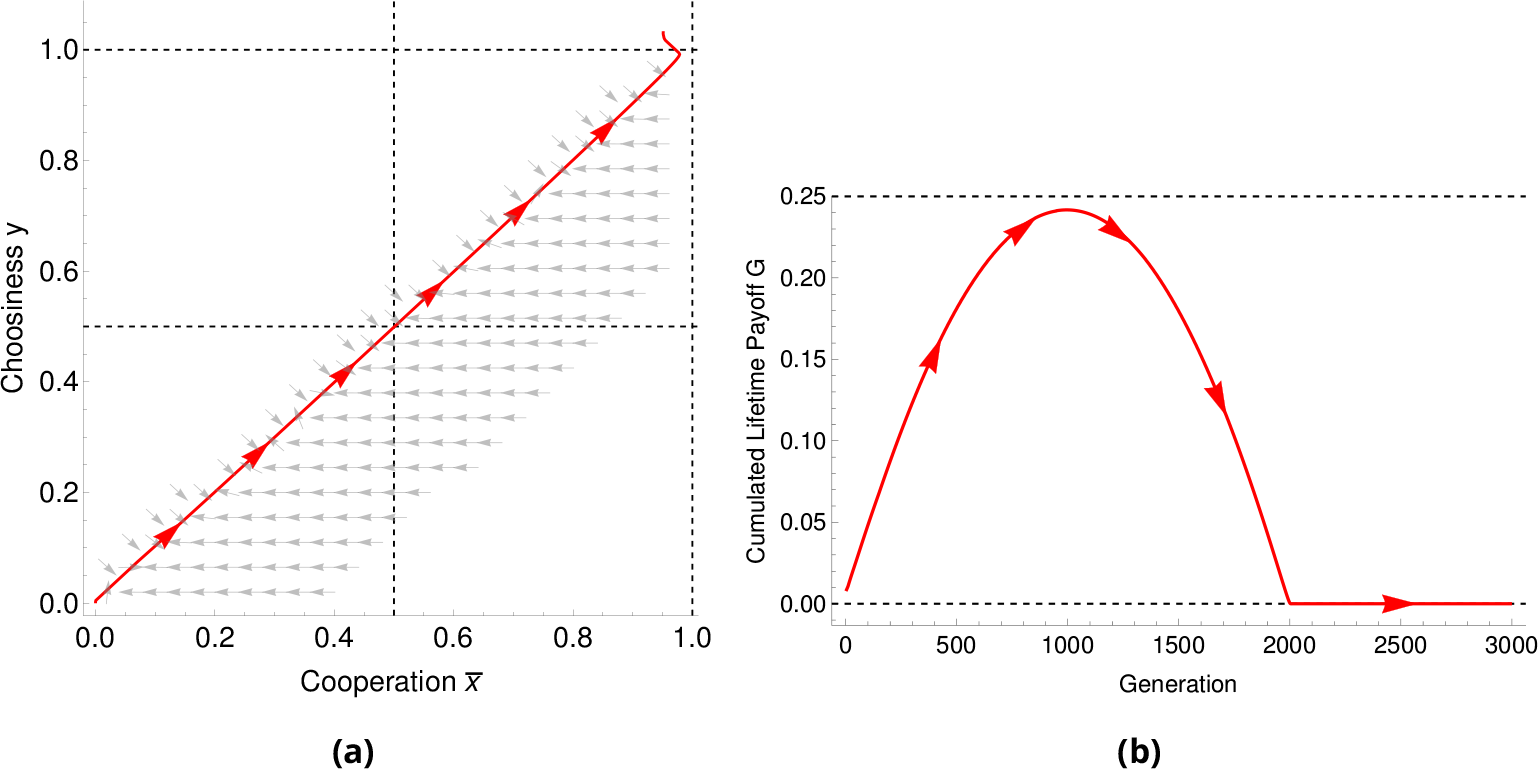
Analytical and numerical results with hard-wired choosiness. **(a)** The grey arrows show the vector field of the selection gradient on both cooperation and choosiness. The red arrows show an evolutionary trajectory starting from an initial selfish population 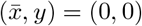. **(b)** The red arrow shows the corresponding evolution of the cumulative lifetime payoff *G* for a resident individual. Parameters are *c* = 1; *σ* = 0.025; *β* = 1; *τ* = 0.01. The socially optimal solution is 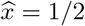 and the interaction becomes profitless if both individuals invest *x* = 1.

This runaway process, however, only occurs if partner choice is very efficient. If partner choice has more frictions, the rise of cooperation and choosiness halts at an intermediate level between 0 and the wasteful threshold. In Figure 2, we plot the level of cooperation (Fig. 2a), the level of choosiness (Fig. 2b) and the average payoff (Fig. 2c) reached at evolutionary stability, in function of the efficiency of partner choice (that is, in function of the parameter *β* controlling the fluidity of the social market and the parameter *σ* controlling the extent of phenotypic variability). As partner choice becomes more efficient, the evolutionarily stable cooperation and choosiness monotonously rise from zero up to the wasteful threshold (Fig. 2a and 2b). Accordingly, the net payoff obtained by individuals at evolutionary stability varies with the efficiency of partner choice in a non-monotonous way. Increasing the efficiency of partner choice has first a positive and then a negative effect on payoff (Fig. 2c). In the extreme, when partner choice is frictionless, cooperation and choosiness increase up to the “wasteful threshold” *x* = 1/*c* at which cooperation is entirely profitless (as was shown in Fig. 1). Note that, in this case, choosiness is even slightly larger than the “wasteful threshold” at equilibrium because, due to phenotypic variability, some individuals cooperate beyond *x* = 1/*c* which makes it adaptive to request higher values of cooperation. In fact, when phenotypic variability is too high (large *σ*), individuals are so choosy at evolutionary equilibrium that the equilibrium level of cooperation is reduced (Fig. 2a). These results have been confirmed in individual-based simulations (see SI).

**Figure 2.**
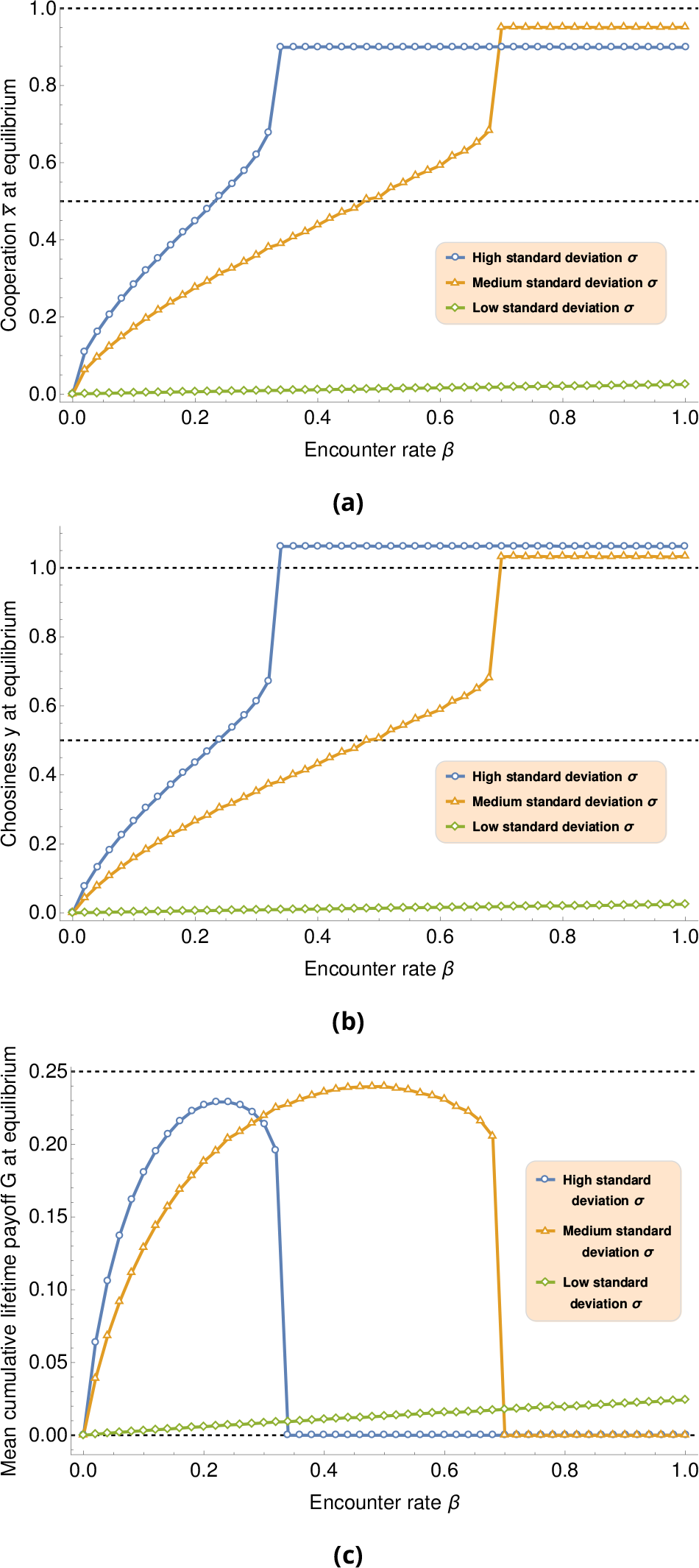
Analytical results for a range of parameters with hard-wired choosiness. Equilibrium values are shown for **(a)** cooperation, **(b)** choosiness and **(c)**cumulative lifetime payoff as a function of the encounter rate *β* to manipulate the market fluidity, and for three values of the standard deviation *σ* = 0.0001; 0.01; 0.02 respectively for low, medium and high phenotypic variability. Other parameters are the same as in Fig. 1

The runaway process can be understood intuitively. In any population, some individuals cooperate more than average, in particular owing to phenotypic variability. As a result, if partner choice is sufficiently fluid, it is adaptive to accept only these hyper-generous partners. Hence, choosiness increases by natural selection beyond the average cooperation level. In turn, this favours individuals who cooperate more than average, i.e. the mean level of cooperation increases by natural selection, etc. The extent to which this process goes on depends, however, on the efficiency of partner choice owing to the existence of a trade-off between the cost and benefit of choosiness. The runaway process stops at the point where the expected benefit of finding a better partner is not worth the risk of remaining alone.

In our model so far, the cost and benefit of switching partner are only determined by two parameters (the market fluidity, *β/τ*, and the amount of phenotypic variability, *σ*). Under more realistic biological assumptions, however, the cost of rejecting a partner should also depend on other parameters. For instance, one could model mortality as a stochastic process. The risk of dying while searching for a new partner would then constitute a supplementary cost of choosiness [72]. In the Supplementary Information, we develop a model based on a Moran process where individuals are subject to a constant mortality rate. As expected, ceteris paribus, the runaway process results in lower levels of cooperation and choosiness at evolutionary equilibrium when the mortality rate is high. Cooperation, however, still rises beyond the socially optimal level, even up to the wasteful threshold, if *β* is large and if the mortality rate is not too high.

Also, in our model, so far, we assume that an individual’s phenotypic level of cooperation is randomized in every encounter. The distribution of cooperative types in the solitary population is thus a fixed and exogenous property. To test the robustness of our results, in the Supplementary Information, we analyse an alternative case where the phenotypic level of cooperation of an individual is randomized only once, at birth. In this case, the distribution of cooperative types in the solitary population is not an exogenous, fixed, property. More cooperative individuals are less likely to be solitary than average because they are rapidly accepted as partners [72]. Hence, the population of solitary individuals tends to be biased toward selfish phenotypes. As a result, the cost of being choosy is larger. Yet, in SI we show that the runaway process still occurs in this case, including up to the “wasteful threshold”, as long as partner choice is efficient enough.

Note that Ferriere *et al.* [49] and Wild and Cojocaru [105] (inspired by Barclay [17]) also showed that partner choice could, under some circumstances, drive the evolution of cooperation up to a “wasteful threshold”. However, in both models, the choosiness strategy was fixed, and not necessarily optimal; it did not evolve jointly with cooperation. The present results are thus more robust and general.

### 3.2 Plastic choosiness

Here, an individual’s choosiness is a reaction norm to her own phenotypic cooperation, and we used search and matching models (see Section 2.3) to derive the two following predictions regarding the evolutionarily stable reaction norm:

i. If the social payoff function is strictly log-supermodular, an individual’s optimal choosiness is a strictly increasing function of her own cooperation (weakly positive assortative matching).
ii. If the market fluidity *β/τ* is high, the reaction norm should be close to *y*\*(*x*) = *x* ∀*x* (strictly positive assortative matching).

We first show that our production function Π is strictly log-supermodular. Indeed, Π(*x*_*i*_, *x*_*j*_)Π(*x*_*k*_, *x*_*l*_) > Π(*x*_*i*_, *x*_*l*_)Π(*x*_*k*_, *x*_*j*_) is equivalent to

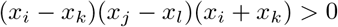

which is true for all *x*_*i*_ > *x*_*k*_ ≥ 0 and *x*_*j*_ > *x*_*l*_. Accordingly, search and matching models show that the optimal choosiness strategy is an increasing reaction norm, i.e. more phenotypically cooperative individuals should also be choosier, leading to a positive assortative matching at equilibrium (phenotypically generous individuals are matched with other generous individuals, and vice versa).

Individual-based simulations confirm this result. Fig. 3 shows the reaction norm at evolutionary equilibrium in these simulations: choosiness is strictly increasing, at least around the levels of phenotypic cooperation that are actually present at equilibrium. Outside this range, selection is very weak on the reaction norm, and we observe larger confidence intervals. As expected, when the market tends to be frictionless, the reaction norm becomes very close to the identity function, that is to a strict positive assortative matching (Fig. 3a and 3b, orange dashed line).

**Figure 3.**
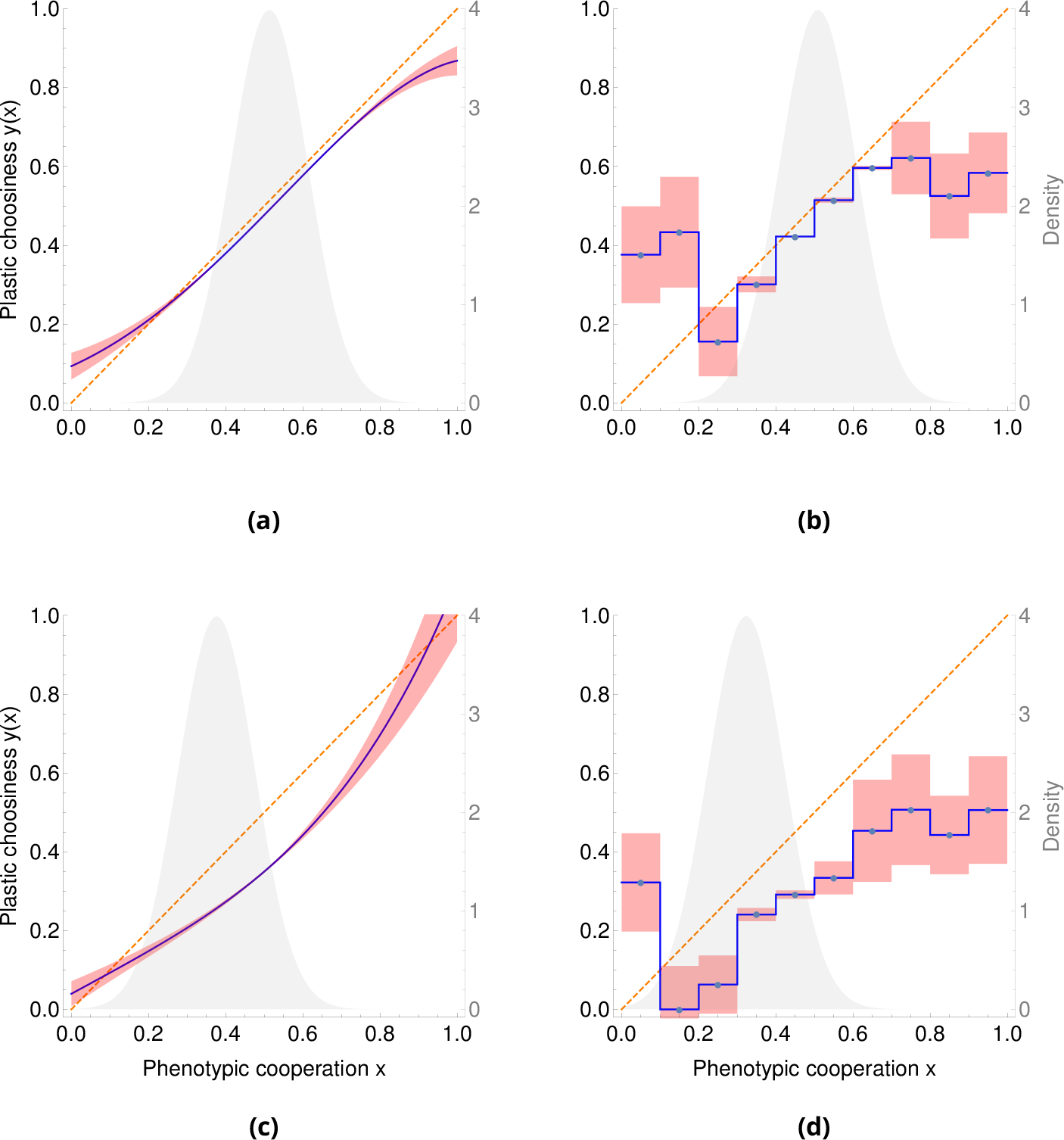
Plastic choosiness at the equilibrium. The equilibrium reaction norms over 30 simulations are shown in blue, and the corresponding 99% confident intervals are shown in red with **(a-b)** high market fluidity *β* = 1, **(c-d)** low market fluidity *β* = 0.01, **(a-c)** a polynomial reaction norm, and **(b-d)** a discrete reaction norm. The orange dashed line is the optimal reaction norm for a frictionless matching market (strong form of positive assortative matching). The distribution of phenotypic cooperation at equilibrium are shown in grey. Parameters are *c* = 1; *σ* = 0.1; *τ* = 0.01; *μ* = 0.001; *σ*_*mut*_ = 0.05; *μ*_*l*_ = 0.05; *N* = 300; *L* = 500.

Importantly, the evolution of a plastic rather than hard-wired choosiness strategy has a key consequence regarding the efficiency of cooperation at evolutionary equilibrium. In contrast with the hard-wired case, when choosiness is plastic cooperation never rises above the socially optimal level. As the efficiency of partner choice (that is, market fluidity) increases, the level of cooperation at evolutionary stability increases but, at most, it reaches the socially optimal level and never more (Fig. 4). In particular, when partner choice is very efficient, cooperation evolves precisely towards the socially optimal level, i.e. the level that maximizes the net total payoff of individuals 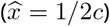.

**Figure 4.**
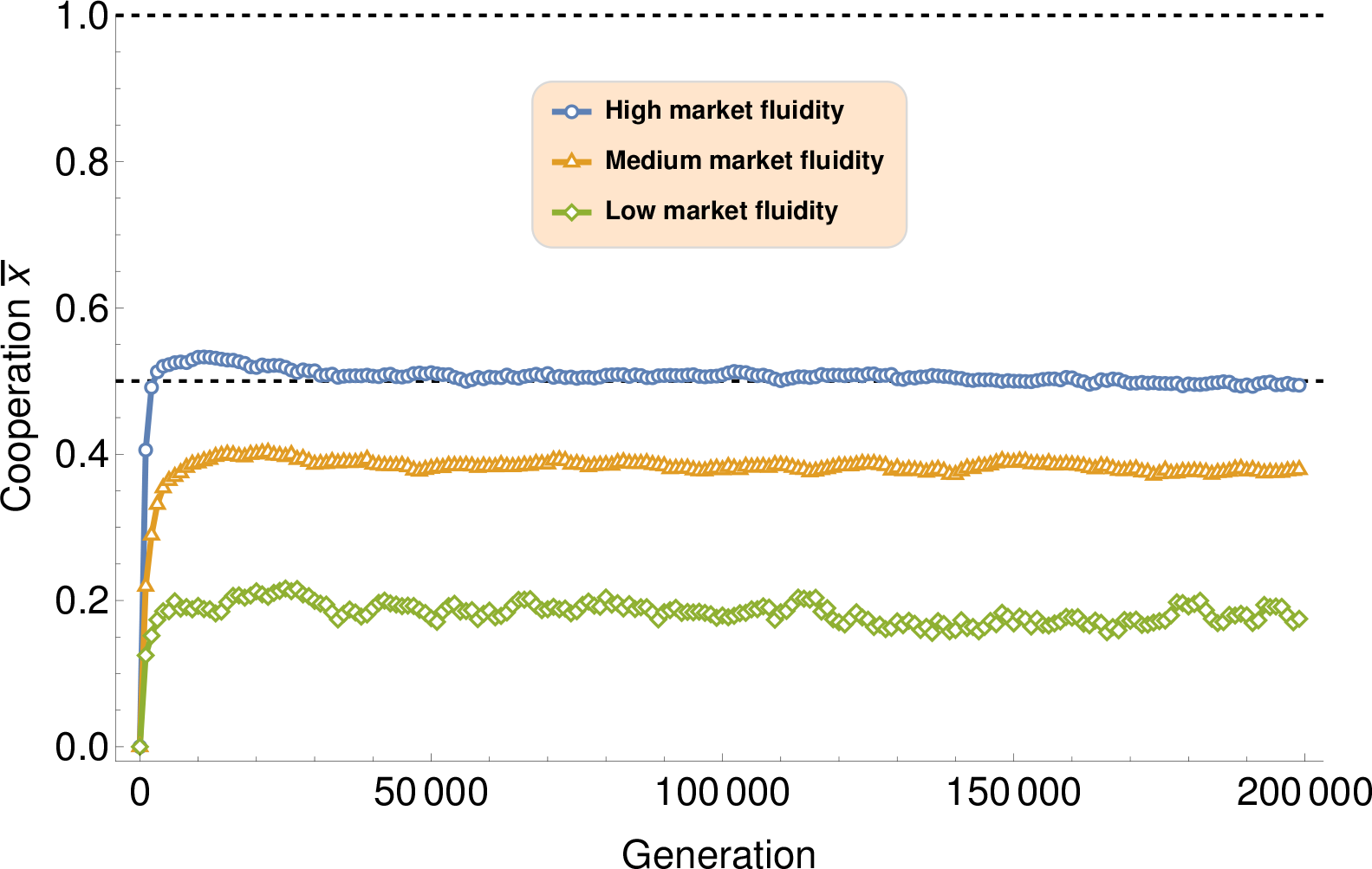
Evolution of cooperation for a polynomial reaction norm. The average cooperation over 30 simulations is shown for three values for the encounter rate *β* = 0.001; 0.01; 0.1 respectively for low, medium and high market fluidity. Other parameters are the same as in Fig. 3. The socially optimal solution is 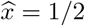 and the interaction becomes profitless if both individuals invest *x* = 1.

This result can also be understood intuitively. In the first model where choosiness was hard-wired, it was adaptive to increase one’s cooperation level beyond the population mean because, by doing so, an individual could switch from “being rejected by everyone”, to “being accepted by everyone”. The runaway process, therefore, proceeded until cooperation had no benefit at all. In contrast, in the present model where choosiness is plastic, increasing one’s cooperation level is beneficial because it allows one to access better partners. Hence, this is useful only provided the benefit of accessing a higher quality partner is larger than the cost of being more cooperative. As a result, cooperation only rises up to the social optimum, where its net benefit is maximized.

## 4 Discussion

Most theoretical works on the evolution of cooperation by partner choice aim at explaining how *some* cooperation can be evolutionarily stable. They do not aim at understanding which specific quantitative *level* of cooperation should evolve. In this paper, we have raised this second question. We have considered a model where cooperation has diminishing returns, such that the most efficient level of cooperation (the level that maximises social welfare) is intermediate. We have investigated whether partner choice can account for the evolution of an efficient level of cooperation in this case. In this aim, we have modelled, both numerically and with individual-based simulations, the joint evolution of two traits: cooperation, the effort invested into helping others, and choosiness, the minimal level of cooperation that an individual is willing to accept in a partner.

In a first model, we have found that the mechanism of partner choice entails no systematic force favouring an efficient level of cooperation. On the contrary, when partner choice is effective enough, the level of cooperation increases evolutionarily toward very large values, beyond the socially optimal level. In the extreme, when partner choice is very effective, cooperation even increases up to a level where its cost entirely cancels out its benefit. That is, at evolutionary equilibrium, individuals gain the same payoff than if they had not cooperated at all.

To understand intuitively, consider a population with a given distribution of cooperation levels, with some particularly generous individuals, some particularly stingy individuals, and a given mean cooperation level. In such a population, provided that the variability of cooperation is sufficiently large and the market sufficiently fluid, it is always adaptive to accept only partners that are slightly better than average [72]. Hence, natural selection favours individuals with a choosiness always slightly larger than the average cooperation level. In turn, this choosiness selects for mutants whose cooperation level is larger than the mean, which leads to a gradual increase in cooperation. Importantly, this runaway process has no particular reason to stop when cooperation is maximally efficient. Rather, it stops when the cost of searching for more generous individuals exceeds the benefit of interacting with them (Fig. 2). As long as partner choice is effective (i.e. the cost of searching is low), it is always worth trying to find a better than average partner, irrespective of whether the current mean level of cooperation is below or beyond the socially optimal level. Hence, partner choice can prompt individuals to invest into counterproductive forms of cooperation to outbid others, leading to an eventually fruitless arms race.

In a second approach, in line with matching models from the economic literature, we have designed a model in which choosiness is implemented as a reaction norm to the individual’s own cooperation level (see Section 2.3), the shape of which evolves by natural selection. In this case, both our analytical model and complementary individual-based simulations show that the evolutionarily stable reaction norm is a monotonously increasing function of cooperation (Fig. 3). This implies that more generous individuals are also choosier, leading to a positive assortative matching: generous individuals tend to interact with other generous individuals, and vice versa. Furthermore, if the biological market is fluid enough (i.e. if the cost of changing partner is low), this positive assortative matching becomes very close to a perfect matching in which individuals with a given level of cooperation always interact with other individuals with the exact same level (Fig. 3a and 3b).

In this case, and in sharp contrast with the model in which choosiness is a hard-wired trait, cooperation does not reach the counterproductive level where its cost cancels out its benefit when partner choice is very cheap (Fig. 4). More precisely, when the market is very fluid, the evolutionarily stable cooperation becomes very close to the social optimum, i.e. the amount of cooperation that maximizes the sum of the partners’ payoffs. This can also be understood intuitively. Because of the strict assortment between cooperative types, individuals with a given cooperation level interact with other individuals with the exact same level. Hence, pairs of individuals become the effective units of selection, like if interactions occurred among genetic clones [1, 4, 48, 106]. Consequently, the socially optimal level of cooperation is favoured.

Hence, the fruitless runaway of cooperation that occurs in a model with hard-wired choosiness is a consequence of the assumption that individuals cannot optimally adapt their degree of choosiness to local circumstances. If individuals are allowed to behave optimally, which entails in the present case to adapt plastically their choosiness to their own generosity, then partner choice looks less like a competition to outbid others, and more like a competition to form efficient partnerships with others, which leads to a very different outcome regarding the net benefits of cooperation.

Previous work has shown that assortative matching favours the evolution of cooperation [25, 48, 58]. For instance, in kin selection, assortment between relatives drives the evolution of cooperation [57, 83]. To our knowledge, Wilson & Dugatkin [106] first discussed the consequences of assortative matching for the evolution of socially efficient levels of cooperation. Alger & Weibull [6, 7] have studied the evolution of social preferences, rather than strategies, under assortative matching. However, both analyses did not explicitly model a partner choice strategy, let alone the evolution of this strategy, but merely assumed that assortment occurs in one way or another. In contrast, here, we have studied the joint evolution of choosiness and cooperation, showing how a positive assortative matching can emerge from a simple partner choice mechanism.

In another related work, using individual-based simulations McNamara *et al.* [72] also observed a form of assortative matching in the joint evolution of cooperation and choosiness. One of the main differences with the present approach, however, is that they assumed that the variability of cooperation is maintained at the genetic level, via a high mutation rate, rather than at the phenotypic level. Under this assumption, negative selection on inefficient mutants (either too choosy or too generous) generates linkage disequilibrium between cooperation and choosiness, resulting in a positive assortative matching. For this reason, their work is more similar to our second model where choosiness is plastic than to our first model where choosiness is hard-wired. In McNamara *et al.*’s simulations [72], however, in contrast with our results, cooperation never reaches the socially optimal level (in the model where they consider a payoff function with diminishing returns). In a complementary analysis (see SI), we showed that this could be a consequence of their assumption that the genetic mutation rate is very high, which prevents natural selection from fully optimizing social strategies.

Some scholars have already imported principles from matching theory into evolutionary biology, especially in the field of sexual selection. Johnstone *et al.* [63] and Bergstrom & Real [24] have used matching models, respectively with and without search frictions, to shed light on mutual mate choice. Both works focused on the evolution of choosiness with a given, fixed distribution of individual’s quality. As we have previously shown, the intensity of assortment may have a dramatic impact on the evolution of the chosen trait (cooperation, in our case). For instance, further models could investigate the precise limits of the runaway processes that occur on weaponry, or on ornamental traits, in sexual selection. More generally, matching models could be helpful to analyse a large variety of biological markets [59, 75, 76], including inter-specific mutualisms, such as mycorrhizal symbiosis or plant-rhizobia relationships [64, 65, 90].

As for the human case in particular, several lines of evidence suggest that partner choice is a likely candidate as a key driving force in the evolution of cooperation. Numerous experimental studies have shown that human beings indeed do choose their social partners in function of their cooperative reputation [16, 19–22, 47, 79, 93, 95, 96, 107]. Anthropological observations show that defection in traditional societies is mostly met with a passive abandon rather than with more defection in return (see [22] for a review). Also, several theoretical studies have shown that partner choice can account for the evolution of other important properties of human cooperation, such as the fact that its benefits are often shared in proportion to everyone’s respective effort in producing them [11, 12, 41, 43–45, 97].

Regarding the quantitative level of cooperation, observations show that humans have precise preferences regarding the amount of effort that shall be put into helping others. Daily life contains ample examples of these preferences. For instance, we hold the door for others in subway stations, but only when they are sufficiently close to the door already, not when they are very far from it. And this is true quite generally. As experiments in real settings demonstrate, we have preferences for specific amounts of cooperation, neither too little, nor too much [67, 85]. Sometimes this preference is expressed in a purely quantitative manner. At other times, the same preference is expressed in a more qualitative way, determining the kinds of cooperative action that we are willing, or unwilling, to perform. In any case, our investment in helping is quantitatively bounded. Moreover, the precise level of effort we are willing to put in cooperation seems to be constrained by considerations of social efficiency. Individuals help one another only when it is mutually advantageous, that is when the cost of helping is less than the benefit of being helped. Additionally, recent evolutionary modellings of risk pooling have revealed the socially optimal nature of helping behaviours [2, 5, 36, 42, 60]. They have shown that people’s systems of mutual help correspond to the most efficient systems of risk pooling in a volatile environment.

In this paper, we have shown that partner choice can foster the evolution of such an intermediate and efficient amount of cooperation, neither too little nor too much. But we have also shown that the precise evolutionarily stable amount of cooperation should depend on the fluidity of the biological market, and can range from a very low level of cooperation, up to the socially optimal level (Fig. 4). A number of anthropological studies suggest that contemporary hunter-gatherer societies exhibit high levels of spatial mobility [22, 71]. Therefore, it seems plausible that biological markets were highly fluid in the social structure that our ancestors experienced. Our model predicts that, in this case, the amount of effort invested into cooperation should become very close to the social optimum. Therefore, partner choice can account for the evolution of human preferences concerning social efficiency.

One could wonder, however, whether other models than partner choice could account for the evolution of a socially optimal level of cooperation as well. The most influential model on the evolution of quantitative cooperation among non-kin is the continuous version of the iterated prisoner’s dilemma [9, 13, 66, 68, 81, 99, 100]. In this game, André & Day [13] have shown that the only evolutionarily stable level of investment is the one that maximises the total benefit of the interaction, i.e. that natural selection does eventually favour the socially optimal amount of cooperation (see also [26, 27, 54, 82] in a discrete version of the iterated prisoner’s dilemma). Yet, in this approach, selection for efficient cooperation is only a second-order force, which plays a significant role only because André & Day [13] assumed the absence of other first-order effects. For instance, a slight cognitive cost of conditional behaviour would have prevented the evolution of efficient cooperation in their model. In another related study, Akçay & Van Cleve [1] have shown that socially optimal cooperation is favoured when individuals play a specific class of behavioural responses to others’ cooperative actions. They have also shown that, for a specific case of their model, these behavioural responses can evolve by natural selection under low levels of relatedness Here, we have shown that, under the effect of partner choice, efficient cooperation is favoured by first-order selective effects even in the total absence of genetic relatedness. This occurs because, unlike reciprocity, partner choice is a *directional* enforcement mechanism. Whereas reciprocity merely stabilizes any given level of cooperation (a principle called the folk theorem, see [14, 31]), partner choice directionally favours the most efficient level.

One limit of our model is that we did not introduce an explicit mechanism for reputation. We simply assumed that, in a way or another, individuals have reliable information regarding the cooperation level of others, but we did not model the way in which they obtain this information. Costly signalling theory proposes that some cooperative behaviours are costly signals of an individual’s quality or willingness to cooperate [10, 18, 28, 29, 56, 69]. Such signals could, in theory, be far from socially efficient [56]. However, further analyses are needed to rigorously model signalling in the context of a biological market.

## Acknowledgements

This work was supported by ANR-10-LABX-0087 IEC and ANR-10-IDEX-0001-02 PSL. This is contribution 2019-004 of the Institut des Sciences de l’Evolution de Montpellier (UMR CNRS 5554). This preprint has been reviewed and recommended by Peer Community In Evolutionary Biology (https://dx.doi.org/10.24072/pci.evolbiol.100063). We thank the Recommender, Erol Akçay, and two anonymous reviewers for very helpful comments on previous versions of this paper.

