## Supporting Information for "Why cooperation is not running away"

### "Hard-wired choosiness" model

We suppose a infinite size population of individuals with lifespan $L$. Solitary individuals randomly encounter each other at a fixed rate $\beta$ (See section *"Linear" and "quadratic" search* for an alternative assumption). If both individuals accept each other, they leave the solitary state and enter an interaction state that dissolves at rate $\tau$. Each individual $i$ is genetically characterized by two traits: her cooperation level $x_i$, and her choosiness $y_i$. Cooperation level $x_i$ represents the quantitative amount of effort that an individual $i$ is willing to invest into cooperation. Choosiness $y_i$ represents the minimal cooperation level that an individual $i$ is willing to accept in a partner.

As in Song and Feldman (2013) or André (2015), we choose to maintain variability in cooperation at the phenotypic level by assuming that individuals do not perfectly control the level they express. The actual investment into cooperation an individual makes follows a truncated to zero normal distribution with mean $\bar{x}$ and standard deviation $\sigma$. The corresponding density function is:

$$f_{\bar{x}}(x) = \begin{cases} \frac{e^{-\frac{(x-\bar{x})^2}{2\sigma^2}}}{\sqrt{2\pi}\sigma\left(1-\frac{1}{2}\text{erfc}\left(\frac{\bar{x}}{\sqrt{2}\sigma}\right)\right)} & x > 0 \\ 0 & x \leq 0 \end{cases}$$

We call cooperation level the genetically encoded cooperation level $\bar{x}_i$ that individuals aim for, and "phenotypic cooperation" the actual level $x_i$ of cooperation that they express after phenotypic noise.

There are two ways to implement this variability. Cooperation can be randomized at birth and stay constant for the whole individual's life, or at every encounter she makes. For the sake of simplicity, we use the first case in our adaptive dynamics model, but we also run agent-based simulations for both cases (see section "Complementary agent-based simulations").

Individuals encounter at random, so their cooperation level are independent. When two individuals $i$ and $j$ encounter, the probability that they are mutually compatible is therefore:

$$\alpha_i = \int_{y_j}^{\infty} \int_{y_i}^{\infty} f_{\bar{x}_i}(x) f_{\bar{x}_j}(y) dx dy$$

We suppose that mutants individuals are rare, and therefore only interact with residents. When in the solitary state, a mutant $m$ gains the solitary payoff normalized to zero per unit of time. When interacting with a resident $r$, she gains a mean social payoff $\bar{\Pi}_m$ per unit of time. $\bar{\Pi}_m$ is the conditional expectation of the social payoff, given that the the mutant and the resident are mutually compatible. Let $MC$ be the event "the mutant and the resident are mutually compatible" and $1_{MC}$ the indicator random variable of the event $MC$.

$$\bar{\Pi}_m = \mathbf{E}[\Pi(x_m, x_r) \mid MC]$$

$$\bar{\Pi}_m = \frac{\mathbf{E}[1_{MC}\Pi(x_m, x_r)]}{P[MC]}$$

$$\bar{\Pi}_m = \frac{\int_{y_r}^{\infty} \int_{y_m}^{\infty} \Pi(x_m, y_r) f_{\bar{x}_m}(x_m) f_{\bar{x}_r}(x_r) dx_m dx_r}{\alpha_m}$$

The mutant fecundity is supposed to be totally explained by her average cumulative lifetime payoff $G_m$. Let $P_S(t)$ be the probability for a mutant to be in the solitary state at time $t$. Accordingly, $P_I(t) = 1 - P_S(t)$ is the probability for a mutant to be involved in an interaction at time $t$. A mutant encounters a mutually compatible partner at rate $\alpha_m\beta$, and any interaction she can be involved in dissolves at rate $\tau$. We can then write the following dynamics:

$$\frac{dP_S(t)}{dt} = \tau P_I(t) - \alpha_m \beta P_S(t)$$

Making the additional assumption that individuals are born solitary ($P_S(0) = 1$), we can integrate the differential equation:

$$P_S(t) = 1 - \frac{\alpha_m \beta}{\alpha_m \beta + \tau}(1 - e^{-(\alpha_m \beta + \tau)t})$$

In the sake of simplicity, we assume that the lifespan of individuals $L$ is always very large in front of the durations of both social interactions and solitary states. Therefore, the fraction of time spent by an individual in the solitary state can be written as $\frac{\tau}{\alpha_m \beta + \tau}$. Now, we can derive the average cumulative lifetime payoff $G_m$:

$$G_m = \bar{\Pi}_m \frac{\alpha_m \beta}{\alpha_m \beta + \tau} L$$

Without loss of generality, we make the assumption that the lifespan of individuals is equal to unity. The cumulative lifetime payoff simplifies to

$$G_m = \frac{\bar{\Pi}_m \alpha_m \beta}{\alpha_m \beta + \tau}$$

### Case without variability ($\sigma = 0$)

Here, every resident individuals have the same cooperation level $\bar{x} = x$ and choosiness $y$. We can study the following three situations.

1. $x < y$ No interaction occurs. Both traits are under genetic drift, therefore situation 1 can sometimes results in situation 3.

2. $x > y$ All interactions occur. Choosiness is under genetic drift. A slightly less cooperative mutant would invade because she will be accepted and gain as much as the residents. So, situation 2 always results in situation 3.

3. $x = y$ All interactions occur. A more cooperative mutant would gain less than the residents. A less cooperative one would be refused by the residents. A choosier mutant would refused the residents. A less choosy mutant could invade by drift, which would lead to situation 2.

We can see that cooperation can never evolve towards higher values when interactions occur. It can only go downward by drift. If one assumes an arbitrarily small cost $\epsilon$ of being choosier, the only stable equilibrium is $(x, y) = (0, 0)$.

### Complementary agent-based simulations

In our first model, in which choosiness is hard-wired, we made the assumption that an individual's cooperation level is randomized at every encounter she makes. We can make the alternative assumption that cooperation is randomized at birth and stay constant for the whole individual's life. In the first case, the proportion of each cooperation level in the solitary population is fixed and is given by the truncated normal distribution with mean $\bar{x}$ and standard deviation $\sigma$. On the other hand, in the second case, the distribution is endogenously determined. Indeed, since they are often accepted as partner, very generous individuals are less likely to be solitary than stingy individuals at any given time. A steady-state distribution is endogenously determined by the individuals' level of choosiness and the market fluidity.

Thus, the cost of switching partner is higher when the cooperation level is randomized at birth than when it is randomized at very encounter. We would expect that this feature "counteracts" the runaway process in the same manner as low values of market fluidity and phenotypic variability (see Figure 2 in the main text). However, agent-based simulations show that both assumptions leads to the same result when the market is arbitrarily fluid enough: the runaway between cooperation and choosiness occurs towards high values (Fig. S1 and S2). At the equilibrium, individuals are so choosy than the only cooperation levels that are accepted are socially inefficient ($x > 1/c$).

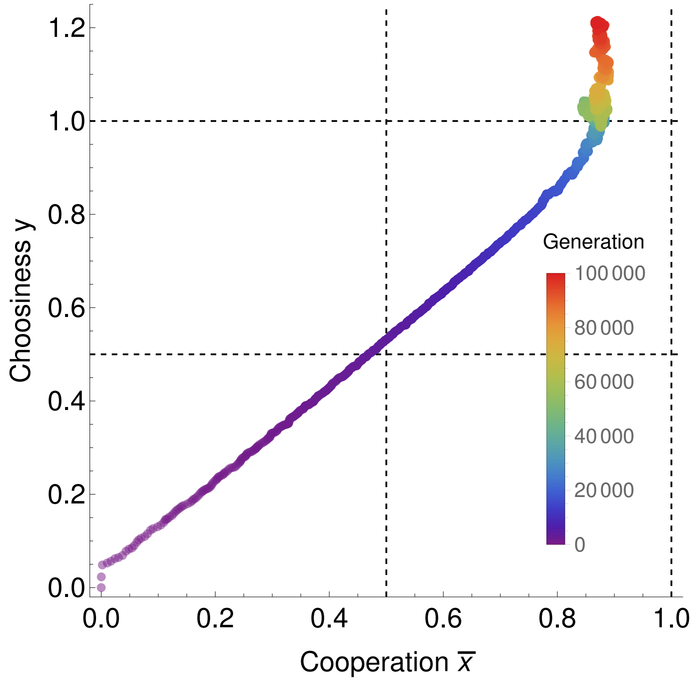


Figure S1: Average trajectory for cooperation and choosiness over 30 simulations with noise at every encounter. Parameters are $\sigma = 0.05$; $\beta = 1$; $\tau = 0.01$; $\mu = 0.001$; $\sigma_{mut} = 0.05$; $N = 300$; $L = 500$. The socially optimal solution is $\widehat{x} = 1/2$ and the interaction becomes profitless if both individuals invest $x = 1$.

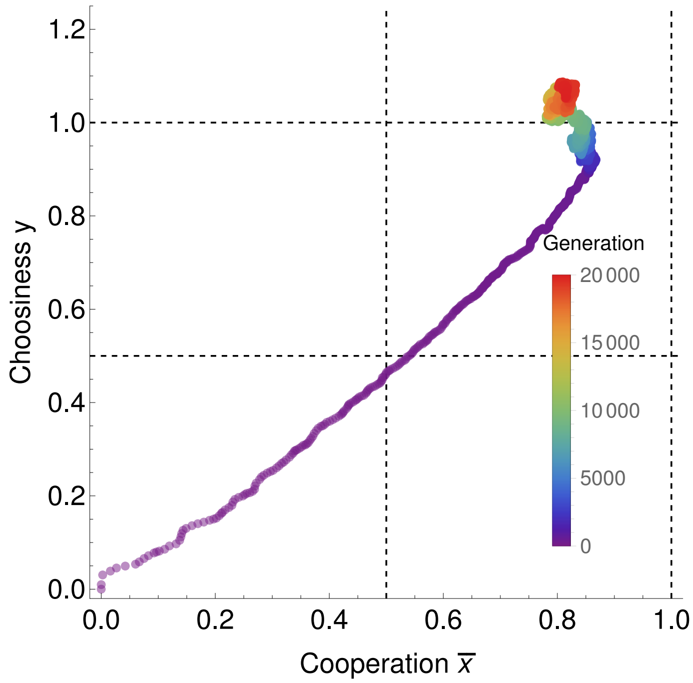


Figure S2: Average trajectory for cooperation and choosiness over 30 simulations with noise at birth. Same parameters as in Fig. S1, except for $\sigma = 0.075$.

### Moran process with a constant mortality rate

Another feature that could counteract the runaway process is the presence of a constant, stochastic probability of dying (this feature is explored in McNamara et al. 2008). Indeed, if an individual rejects a partner, she "runs the risk" to die during her solitary state, which would be a waste of time from a fitness perspective. In our adaptive dynamics model as well as in all of our agent-based simulations, we have used a Wright-Fisher model with non-overlapping generations and where individuals' lifespan is fixed and is very large in front of the duration of both paired and solitary states. A more realistic assumption would be that individuals randomly die according to a mortality rate. In this section, we analyse a Moran process (with overlapping generations) using agent-based simulations. Each individual dies at rate $m$. When an individual dies, she is replaced by another individual whose genotype is chosen according to each individual's fecundity.

Figure S3 shows the equilibrium values of cooperation and choosiness at the evolutionary equilibrium for a very fluid market. When the mortality rate is low, the runaway process goes up to the "wasteful threshold" as in our previous result (Fig. S1). However, when the mortality rate is high, the runaway stops at intermediate values of cooperation and choosiness. In other words, mortality plays a similar role as market fluidity: it is a component of the cost of switching partner. Therefore, it shapes the evolutionary pressure acting on the evolution of choosiness and cooperation.

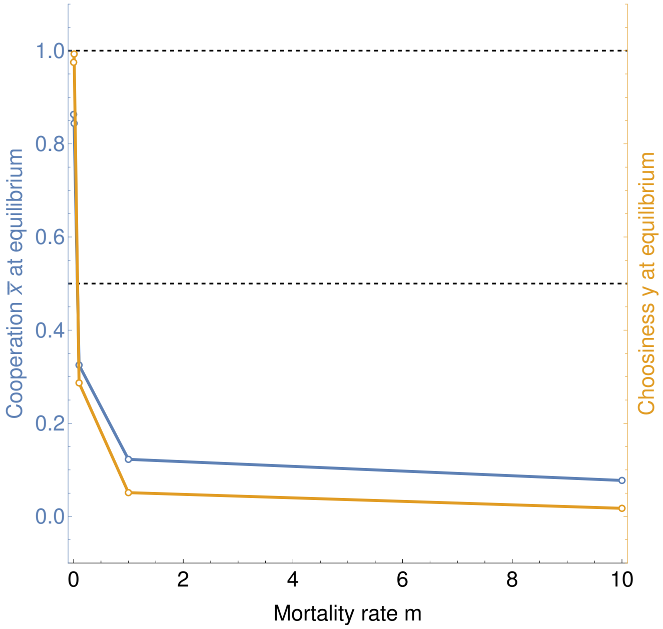


Figure S3: Mean equilibrium value of the cooperation level and the choosiness level as a function of the mortality rate (30 simulations for each point). Same parameters as in Fig. S1 (very fluid market).

### "Linear" and "quadratic" search

In a seminal paper on the modelling of markets with search friction, Diamond and Maskin (1979) identified two assumptions concerning the search process. The search is said to be "linear" if the probability for an individual to find a solitary partner is fixed. On the other hand, the search is said to be "quadratic" if this probability increases with the proportion of solitary individuals. In this case, the paired individuals interfere with the encounters of solitary individuals. This is similar to the mass-action kinetics that is used in epidemiology to model encounters between susceptible and infectious individuals.

For the sake of simplicity, we have used so far a "linear" search by assuming that the encounter rate $\beta$ is fixed. We run complementary simulations of our first model (hard-wired choosiness) and we show that our result is robust to the search assumption used (Fig. S4).

Under the "quadratic" search assumption, the encounter rate is $\beta = \eta \times P_S$, with $\eta$ being a constant.

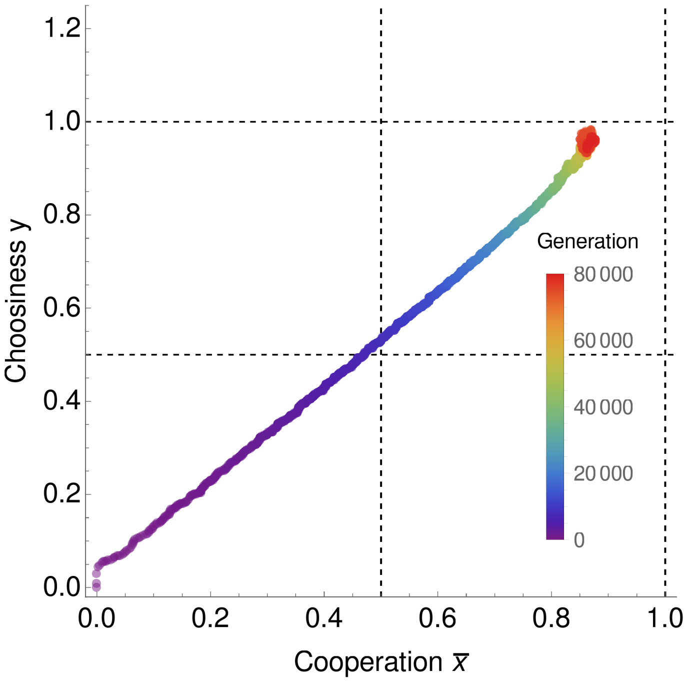


Figure S4: Average trajectory for cooperation and choosiness over 30 simulations with quadratic search. Same parameters as in Fig. S1, except for $\eta = 1$.

The analysis of the market fluidity differs between the "linear" and "quadratic" search. With the "linear" search, when $\beta$ is high (and $\tau$ is low), individuals should be choosy because the mean time before finding a partner is short (and the mean interaction time is long). With the "quadratic" search, the effect of $\eta$ and $\tau$ is more complex. Both $\eta$ and $\tau$ have an effect on the proportion $P_S$ of solitary individuals in the population. Indeed, when $\eta$ is high and $\tau$ is low, more individuals are paired at any given time, hence the encounter rate $\beta = \eta \times P_S$ is reduced.

As in the *"Hard-wired choosiness" model* section, we can derive the proportion $P_S$ of solitary individuals in the population under "quadratic" search. If we assume that the resident individuals have a probability $\alpha$ of making a mutually compatible encounter:

$$\frac{dP_S(t)}{dt} = \tau P_I(t) - \alpha\beta P_S(t)$$

$$\frac{dP_S(t)}{dt} = \tau P_I(t) - \alpha\eta {P_S}^2(t)$$

With the additional assumption that individuals are born solitary ($P_S(0) = 1$):

$$P_S(t) = \frac{\sqrt{\tau}\sqrt{4\alpha\eta+\tau}\tanh\left(\tanh^{-1}\left(\frac{2\alpha\eta+\tau}{\sqrt{\tau}\sqrt{4\alpha\eta+\tau}}\right) + \frac{1}{2}t\sqrt{\tau}\sqrt{4\alpha\eta+\tau}\right) - \tau}{2\alpha\eta}$$

Assuming that the individual's lifespan $L$ is very large, the proportion of solitary individuals stabilizes at $\frac{2\tau}{\sqrt{\tau(4\alpha\eta+\tau)}+\tau}$.

The effective encounter rate of a solitary individual is therefore $\beta = \eta \times \frac{2\tau}{\sqrt{\tau(4\alpha\eta+\tau)}+\tau}$ which is an increasing function of both $\eta$ and $\tau$. Thus, even though the proportion of solitary individuals decreases when $\eta$ is large, overall, the market fluidity increases. The effect of $\tau$ is twofold. When $\tau$ is low, individuals interact for a long time, so the market fluidity increases. Yet, when $\tau$ is low, the encounter rate $\beta$ shrinks, which decreases the market fluidity. In short, under "quadratic" search, $\eta$ and $\tau$ determinate the market fluidity, but the splitting rate $\tau$ has a more complex effect than in the "linear" case.

### High mutation rate - Comparison with McNamara et al. (2008)

McNamara et al. (2008) studied the joint evolution of cooperation and choosiness. They implemented variability at the genetic level by assuming a high mutation rate. Negative selection on inefficient mutants (either too choosy or too generous) creates a linkage disequilibrium between the two traits, resulting in a form of positive assortative matching. We can, therefore, compare their approach with our second model where choosiness is plastic, since assortative matching cannot take place in our first model where choosiness is hard-wired.

By means of individual-based simulations, McNamara et al. (2008) showed that cooperation stabilizes at intermediate levels when a payoff function with diminishing returns is used. However, cooperation never reaches the socially optimal level, no matter how cheap switching partner is, or how much variability is assumed. We run complementary simulations to confirm this result in our biological market framework varying the encounter rate $\beta$, mutation rate $\mu$ and the mutation standard deviation $\sigma_{mut}$ (Fig S5 and S6).

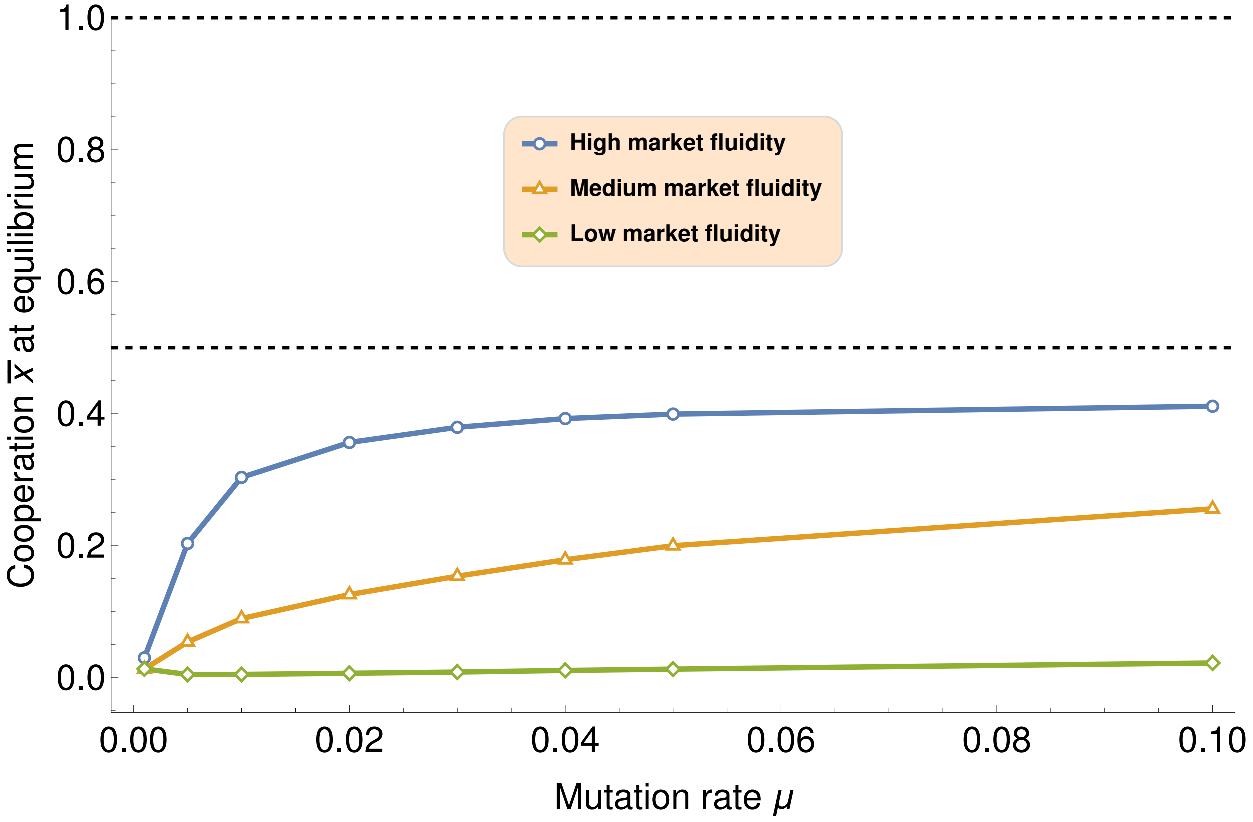


Figure S5: Mean equilibrium value of the cooperation level as a function of the mutation rate $\mu$ and three values of the encounter rate $\beta = 0.001; 0.1; 1$ respectively for low, medium and high market fluidity (30 simulations for each parameter combination). Parameters are $\sigma = 0$; $c = 1$; $\tau = 0.01$; $\sigma_{mut} = 0.05$; $N = 300$; $L = 500$. The socially optimal solution is $\widehat{x} = 1/2$.

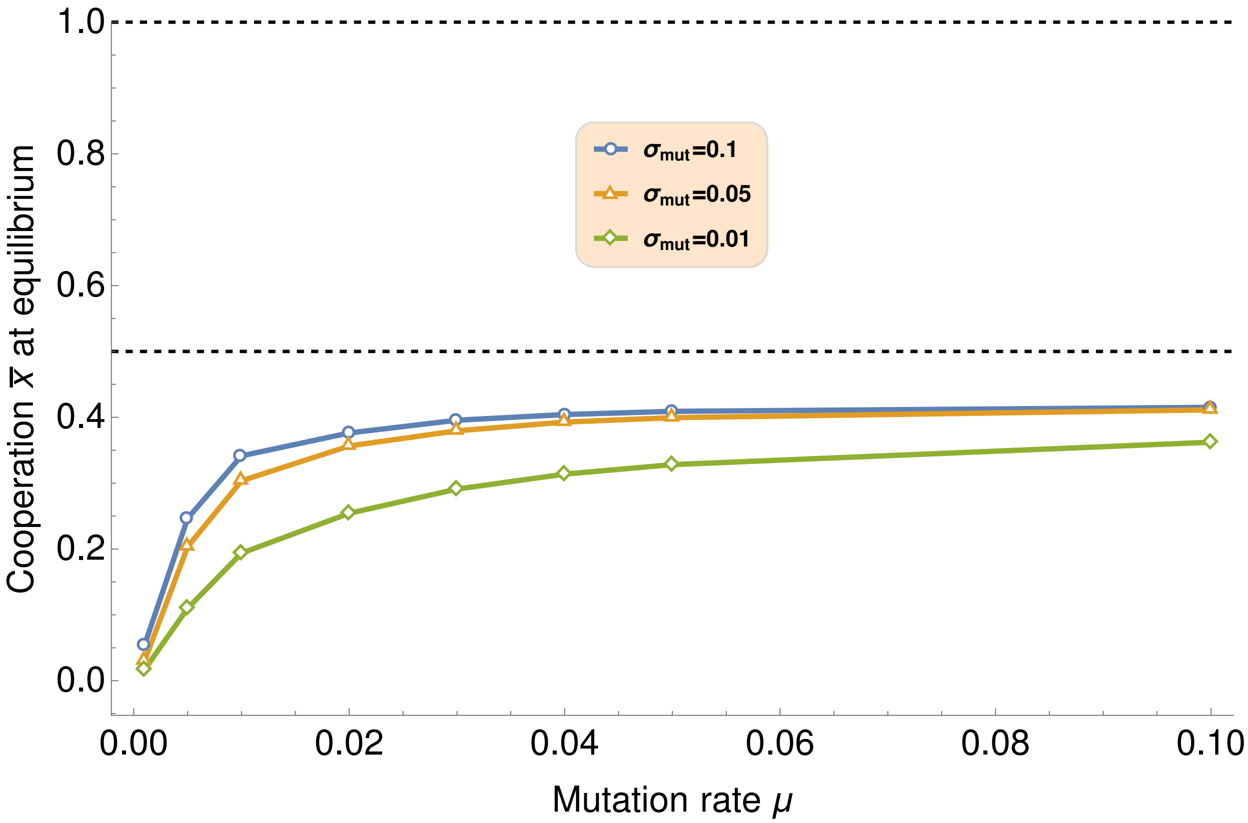


Figure S6: Mean equilibrium value of the cooperation level as a function of the mutation rate $\mu$ and three values of the standard deviation for mutation $\sigma_{mut} = 0.01; 0.05; 0.1$ (30 simulations for each parameter combination). Same parameters as in Fig. S5, except for $\beta = 1$.

The reason why cooperation never reaches the socially optimal level might be the impossibility for all individuals to behave optimally. Indeed, negative selection acts on maladapted individuals, but the mutation process always generates new ones. Clearly this process cannot generate a strict positive assortative matching (where likes interact exactly with likes). Fig S7 shows a comparison of the correlations between the two traits for two different assumptions: a) high mutation rate, and b) phenotypic noise and polynomial reaction norm (as in our second model). Both regression lines indicate an assortment very close to the strictly positive assortative matching. Yet, the genetic variability assumption yields a much higher standard error of the estimate $\sigma_{est}$, indicating that a large proportion of the individuals deviates from the optimal strategy.

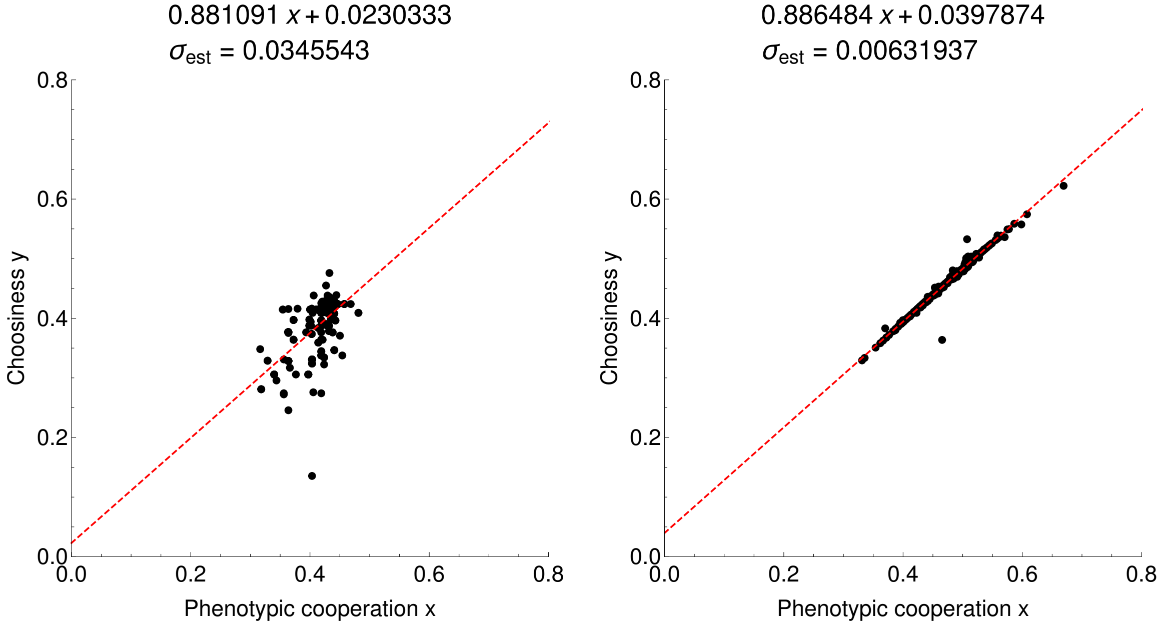


(a) (b)

Figure S7: Correlation between individuals' cooperation and choosiness for two particular simulations. Linear regression coefficients are indicated, as well as the standard error of the estimate $\sigma_{est}$. (a) High level of genetic variability $\mu = 0.05$. (b) High phenotypic variability $\sigma = 0.05$ and plastic choosiness (polynomial function of the cooperation level). Parameters are $\beta = 1$; $\tau = 0.01$; $\sigma_{mut} = 0.05$; $N = 300$; $L = 500$.

The cooperation level stabilizes at the socially optimal level only if the positive assortative matching is strong enough. This cannot be the case if too many maladapted individuals are generated by the mutation process. Therefore, the presence of individuals that deviate from optimality, especially mutants who are too generous mutants and not choosy enough, prevents the evolution of the most socially efficient level of investment in cooperation.
